# Rapid evolutionary diversification of the *flamenco* locus across simulans clade *Drosophila* species

**DOI:** 10.1101/2022.09.29.510127

**Authors:** Sarah Signor, Jeffrey Vedanayagam, Bernard Y. Kim, Filip Wierzbicki, Robert Kofler, Eric C. Lai

## Abstract

Effective suppression of transposable elements (TEs) is paramount to maintain genomic integrity and organismal fitness. In *D. melanogaster*, *flamenco* is a master suppressor of TEs, preventing their movement from somatic ovarian support cells to the germline. It is transcribed by Pol II as a long (100s of kb), single-stranded, primary transcript, that is metabolized into Piwi-interacting RNAs (piRNAs) that target active TEs via antisense complementarity. *flamenco* is thought to operate as a trap, owing to its high content of recent horizontally transferred TEs that are enriched in antisense orientation. Using newly-generated long read genome data, which is critical for accurate assembly of repetitive sequences, we find that *flamenco* has undergone radical transformations in sequence content and even copy number across *simulans* clade Drosophilid species. *D. simulans flamenco* has duplicated and diverged, and neither copy exhibits synteny with *D. melanogaster* beyond the core promoter. Moreover, *flamenco* organization is highly variable across *D. simulans* individuals. Next, we find that *D. simulans* and *D. mauritiana flamenco* display signatures of a dual-stranded cluster, with ping-pong signals in the testis and/or embryo. This is accompanied by increased copy numbers of germline TEs, consistent with these regions operating as functional dual stranded clusters. Overall, the physical and functional diversity of *flamenco* orthologs is testament to the extremely dynamic consequences of TE arms races on genome organization, not only amongst highly related species, but even amongst individuals.

## Introduction

*Drosophila* gonads exemplify two important fronts in the conflict between transposable elements (TEs) and the host – the germline (which directly generates gametes), and somatic support cells (from which TEs can invade the germline) (1, 2). The strategies by which TEs are suppressed in these settings are distinct (3), but share their utilization of piwi-interacting RNAs (piRNAs). These are ~24-32 nt RNAs that are bound by the PIWI subclass of Argonaute proteins, and guide them and associated cofactors to targets for transcriptional and/or post-transcriptional silencing (4–7).

Mature piRNAs are processed from non-coding piRNA cluster transcripts, which derive from genomic regions that are densely populated with TE sequences (7–9). However, the mechanisms of piRNA biogenesis differ between gonadal cell types. In the germline, piRNA clusters are transcribed from both DNA strands through non-canonical Pol II activity (6, 10–12), which is initiated by chromatin marks rather than specific core promoter motifs. Moreover, co-transcriptional processes such as splicing and polyadenylation are suppressed within dual strand piRNA clusters (13, 14). On the other hand, in ovarian somatic support cells, piRNA clusters are transcribed from a typical promoter as a single stranded transcript, which can be alternatively spliced as with protein-coding mRNAs (15–18). These rules derive in large part from the study of model piRNA clusters (i.e. the germline *42AB* and somatic *flamenco* piRNA clusters). For both types, their capacity to repress invading TEs is thought to result from random integration of new transposons into the cluster (19). As such, piRNA clusters are adaptive loci that play central roles in the conflict between hosts and TEs.

The location and activity of germline piRNA clusters are stochastic and evolutionarily dynamic, as there are many copies of TE families in different locations that may produce piRNAs (9, 20). By contrast, somatic piRNA clusters are not redundant and a single insertion of a TE into a somatic piRNA cluster should be sufficient to prevent that TE from further transposition (1, 18). Thus, *flamenco* should contain only one copy per TE family (18), which is true in the *flamenco* locus of *D. melanogaster* (18). *flamenco* is also the only piRNA cluster which produces a phenotypic effect when altered, as germline clusters can be deleted with no consequences (9).

*flamenco* has been a favored model for understanding the piRNA pathway since the discovery of piRNA mediated silencing of transposable elements (6). *flamenco* spans >180 kb of repetitive sequences located in *β*-heterochromatin of the X chromosome (21). Of note, *flamenco* was initially identified, prior to the formal recognition of piRNAs, via transposon insertions that de-repress *gypsy*, *ZAM*, and *Idefix* class elements (21–25). These mutant alleles disrupt the *flamenco* promoter, and consequently abrogate transcription and piRNA production from this locus. By contrast, the recent deletion of multiple model germline piRNA clusters, which eliminate the biogenesis of a bulk of cognate piRNAs, did not de-repress their cognate TEs (9). Thus, the analysis of *flamenco* evolution is presumably more consequential for TE dynamics. Analysis of *flamenco* in various strains of *D. melanogaster* supports that this locus traps horizontally derived TEs to achieve silencing of newly invaded TEs (18). The *flamenco* locus exhibits synteny across the *D. melanogaster* sub-group (26); however, the sequence composition of *flamenco* outside *D. melanogaster* has not been well-characterized (27).

In this study, we compare the *flamenco* locus across 10 strains of simulans-clade species, namely *D. simulans*, *D. mauritiana*, and *D. sechellia*. Analysis of piRNAs from ovaries of five genotypes of *D. simulans* found that *flamenco* is duplicated in *D. simulans*. This duplication is old enough that there is no sequence synteny across copies, even though their core promoter regions and the adjacent *dip1* gene duplications are conserved. *flamenco* has also been colonized by abundant (>40) copies of *R1*, a TE that was thought to insert only at ribosomal genes, and to evolve at the same rate as nuclear genes (28). Furthermore, between different genotypes, up to 63% of TE insertions are not shared within any given copy of *flamenco*. Despite this, several full length TEs are shared between all genotypes in a similar sequence context. This incredible diversity at the *flamenco* locus, even within a single species, suggests there may be considerable variation in its ability to suppress transposable elements across individuals.

Cross-species comparisons further indicate that functions of *flamenco* have diversified. Data from *D. sechellia* and *D. melanogaster* conform with the current understanding of *flamenco* as a uni-strand cluster. However, we find evidence that *D. simulans* and *D. mauritiana flamenco* can act as a dual strand cluster in testis (*D. mauritiana*) and embryos (*D. mauritiana* and *D. simulans*), yielding piRNAs from both strands with a ping pong signal. Overall, we infer that the rapid evolution of *flamenco* alleles across individuals and species reflects highly adaptive functions and dynamic biogenesis capacities.

## Materials and Methods

### Fly strains

The four *D. simulans* lines *SZ232*, *SZ45*, *SZ244*, and *SZ129* were collected in California from the Zuma Organic Orchard in Los Angeles, CA on two consecutive weekends of February 2012 (29–33). *LNP-15-062* was collected in Zambia at the Luwangwa National Park by D. Matute and provided to us by J. Saltz (J. Saltz pers. comm., (34, 35)). *MD251*, *MD242*, *NS137*, and *NS40* were collected in Madagascar and Kenya (respectively) and are described in (36). The *D. simulans* strain *wxD^1^* was originally collected by M. Green, likely in California, but its provenance has been lost (pers. comm. Jerry Coyne). *D. mauritiana (w12)* and *D. sechellia (Rob3c/Tucson 14021-0248.25)* are described in (37).

### Long read DNA sequencing and assembly

*MD242*, four SZ lines and *LNP-15-062* were sequenced on a MinION platform at North Dakota State University (Oxford Nanopore Technologies (ONT), Oxford, GB), with base-calling using guppy (v4.4.2). *MD242*, the four SZ lines, and *LNP-15-062* were assembled with Canu (v2.1) (38) and two rounds of polishing with Racon (v1.4.3) (39). The CA strains were additionally polished with short reads using Pilon (v1.23) (40)(SRR3585779, SRR3585440, SRR3585480, SRR3585391) (29). The first *wxD^1-1^* assembly is described here (41). *MD251*, *NS137*, *NS40* and *wxD^1-2^* were sequenced on a MinION platform at Stanford University. They were assembled with Flye (42), and polished with a round of Medaka followed by a round of pilon (40). Following this contaminants were removed with blobtools (https://zenodo.org/record/845347, (43)), soft masked with RepeatModeler and Repeatmasker (44, 45), then aligned to the *wxD^1^* as a reference with Progressive Cactus (46). The assemblies were finished with reference based scaffolding using Ragout (47). *D. mauritiana* and *D. sechellia* were sequenced with PacBio and assembled with FALCON using default parameters (https://github.com/PacificBiosciences/FALCON)(37).The *D. melanogaster* assembly is described here (48). A summary of the assembly statistics is available in Supplementary Table 1. The quality of cluster assembly was evaluated using the coverage and soft clip quality as described in (20, 49) (Supplementary File 1).

### Short read sequencing and mapping

Short read sequencing was performed by Beijing Genomics Institute on approximately 50 dissected ovaries from adult female flies (*SZ45*, *SZ129*, *SZ232*, *SZ244, LNP-15-062*). Short read libraries from 0-2 hour embryos were prepared from *D. melanogaster*, *wxD^1-2^*, *D. sechellia*, and *D. mauritiana* (SRAXXX) (50). Small RNA from testis is described in (51, 52). *D. melanogaster* OSC small RNA libraries were downloaded from the SRA (SRR11999160). Libraries were filtered for adapter contamination and short reads between 23-29 bp were retained for mapping with fastp (53). The RNA was then mapped to their respective genomes using bowtie (v1.2.3) and the following parameters (-q -v 1 -p 1 -S -a - m 50 --best --strata) (54, 55). The resulting bam files were processed using samtools (56). To obtain unique reads the bam files were filtered for reads with 1 mapping position. To obtain counts files with weighted mapping the bam files were processed using Rsubreads and the featureCounts function (57).

### Defining and annotating piRNA clusters

piRNA clusters were initially defined using proTRAC (58). piRNA clusters were predicted with a minimum cluster size of 1 kb (option “-clsize 1000”), a p-value for minimum read density of 0.07 (option “-pdens 0.07”), a minimum fraction of normalized reads that have 1T (1U) or 10A of 0.33 (option “-1Tor10A 0.33”) and rejecting loci if the top 1% of reads account for more than 90% of the normalized piRNA cluster read counts (option “-distr 1-90”), and a minimal fraction of hits on the main strand of 0.25 (option “-clstrand 0.25”). Note that this ties the piRNA clusters to their function such that participation in the ping pong pathway can be inferred from these patterns. Clusters were annotated using RepeatMasker (v. 4.0.7) and the TE libraries described in Chakraborty et al. (2019) (41, 44). The position of *flamenco* was also evaluated based off of the position of the putative promoter, the *dip1* gene, and the enrichment of *gypsy* elements (15). Fragmented annotations were merged to form TE copies with onecodetofindthemall (59). Fragmented annotations were also manually curated within *flamenco*, particularly because TEs not present in the reference library often have their LTRs and internal sequences classified as different elements.

### Aligning the flamenco promoter region

The region around the *flamenco* promotor was extracted from each genotype and species with bedtools getfasta (60). Sequences were aligned with clustal-omega and converted to nexus format (61). Trees were built using a GTR substitution model and gamma distributed rate variation across sites (62). Markov chain monte carlo iterations were run until the standard deviation of split frequencies was below .01, around one million generations. The consensus trees were generated using sumt conformat=simple. The resulting trees were displayed with the R package ape (63).

### Detecting ping pong signals in the small RNA data

Ping pong signals were detected using pingpongpro (64) This program detects the presence of RNA molecules that are offset by 10 nt, such that stacks of piRNA overlap by the first 10 nt from the 5’ end. These stacks are a hallmark of piRNA mediated transposon silencing. The algorithm also takes into account local coverage and the presence of an adenine at the 10^th^ position. The output includes a z-score between 0 and 1, the higher the z-score the more differentiated the ping pong stacks are from random local stacks.

### Annotating shared and unique TE insertions

To align the TE annotations of homologous piRNA clusters, we first extracted the sequences of the clusters and annotated TEs in these sequences using RepeatMasker (open-4.0.7) with a custom TE library and the parameters: -s (sensitive search), -nolow (disable masking of low complexity sequences), and -no_is (skip check for bacterial IS) (37, 65, 66). Finally, we aligned the resulting repeat annotations with Manna using the parameters -gap 0.09 (gap penalty), -mm 0.1 (mismatch penalty) -match 0.2 (match score) (20, 37). Manna can be used for aligning the annotations of the transposable elements by relying on synteny to determine insertion homology. Alignments were manually checked for inconsistencies arising from assignment to similar TEs (i.e. *gypsy-3* versus *gypsy-5*). TEs were considered to be full length if they were present in at least 70% of their reference length and contained internal sequence as well as two LTRs if applicable.

## Results

### flamenco in the D. simulans clade

We identified *D. simulans flamenco* from several lines of evidence: piRNA cluster calls from proTRAC, its location adjacent to divergently transcribed *dip1*, the existence of conserved core *flamenco* promoter sequences, and enrichment of *gypsy* elements (Figure 1 & 2); Supplementary Table 2). The *flamenco* locus is at least 376 kb in *D. simulans*. This is an expansion compared with *D. melanogaster*, where *flamenco* is only 156 kb (*Canton-S*). In *D. sechellia flamenco* is 363 kb, however in *D. mauritiana* the locus has expanded to at least 840 kb (Supplementary Table 2). This is a large expansion, and it is possible that the entire region does not act as a region controlling somatic TEs. However, evidence that is does include uniquely mapping piRNAs that are found throughout the region and *gypsy* enrichment consistent with a *flamenco*-like locus (Supplementary Figure 1). There are no protein coding genes within the region, and while the neighboring genes on the downstream side of *flamenco* in *D. melanogaster* have moved in *D. mauritiana* (*CG40813*- *CG41562* at 21.5 MB), the following group of genes beginning with *CG14621* is present and flanks *flamenco* as it is annotated. Thus in *D. melanogaster* the borders of *flamenco* are flanked by *dip1* upstream and *CG40813* downstream, while in *D. mauritiana* they are *dip1* upstream and *CG14621* downstream. Between all species the *flamenco* promoter and surrounding region, including the *dip1* gene, are alignable and conserved (Figure 1D).

**Figure 1.**
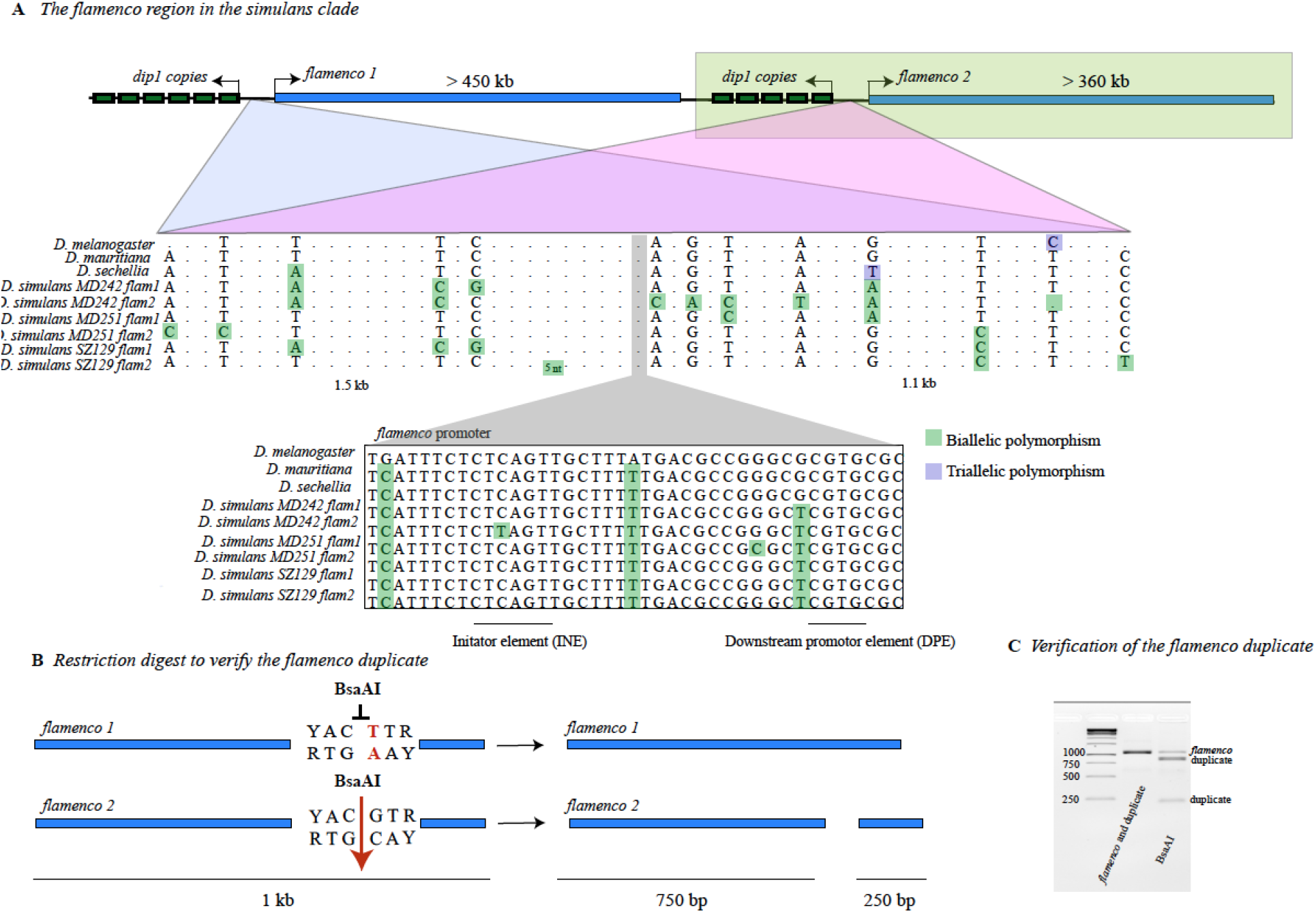
A) The duplication of *flamenco* in the *D. simulans*. Both copies are flanked by copies of the *dip1* gene and copies of the putative *flamenco* promoter. The top portion of the alignment shows ~ 2 kb around the promoter. SNPs are shown if they differentiate copies of *flamenco* within a single genotype of *D. simulans*. Dots do not indicate a single nucleotide, but rather a sequence region where no SNPs differentiate the two copies of *flamenco* within a single genotype. The lower portion illustrates the promoter region with all SNPs illustrated in *D. melanogaster*, *D. sechellia*, *D. mauritiana*, and *D. simulans*. B) A schematic of the restriction digest used to verify the duplicate of *flamenco*. The targeted region is a 1 kb fragment adjacent to the promotor of *flamenco*. Within this region the original *flamenco* copy does not contain a YACGTR site and is not cut by the restriction enzyme BsaAI. The duplicate of *flamenco* is cut into two pieces (750 bp and 250 bp). C) A gel showing the fragments of the original and duplicated copy of *flamenco* before and after digestion with BsaAI. Both copies of *flamenco* are amplified by the primers, in column two of the gel (Supplemental File 2). In column three of the gel, the original copy of *flamenco* is uncut (band 1), while the duplicate of *flamenco* forms two bands at 750 bp (band 2) and 250 bp (band 3).

**Figure 2.**
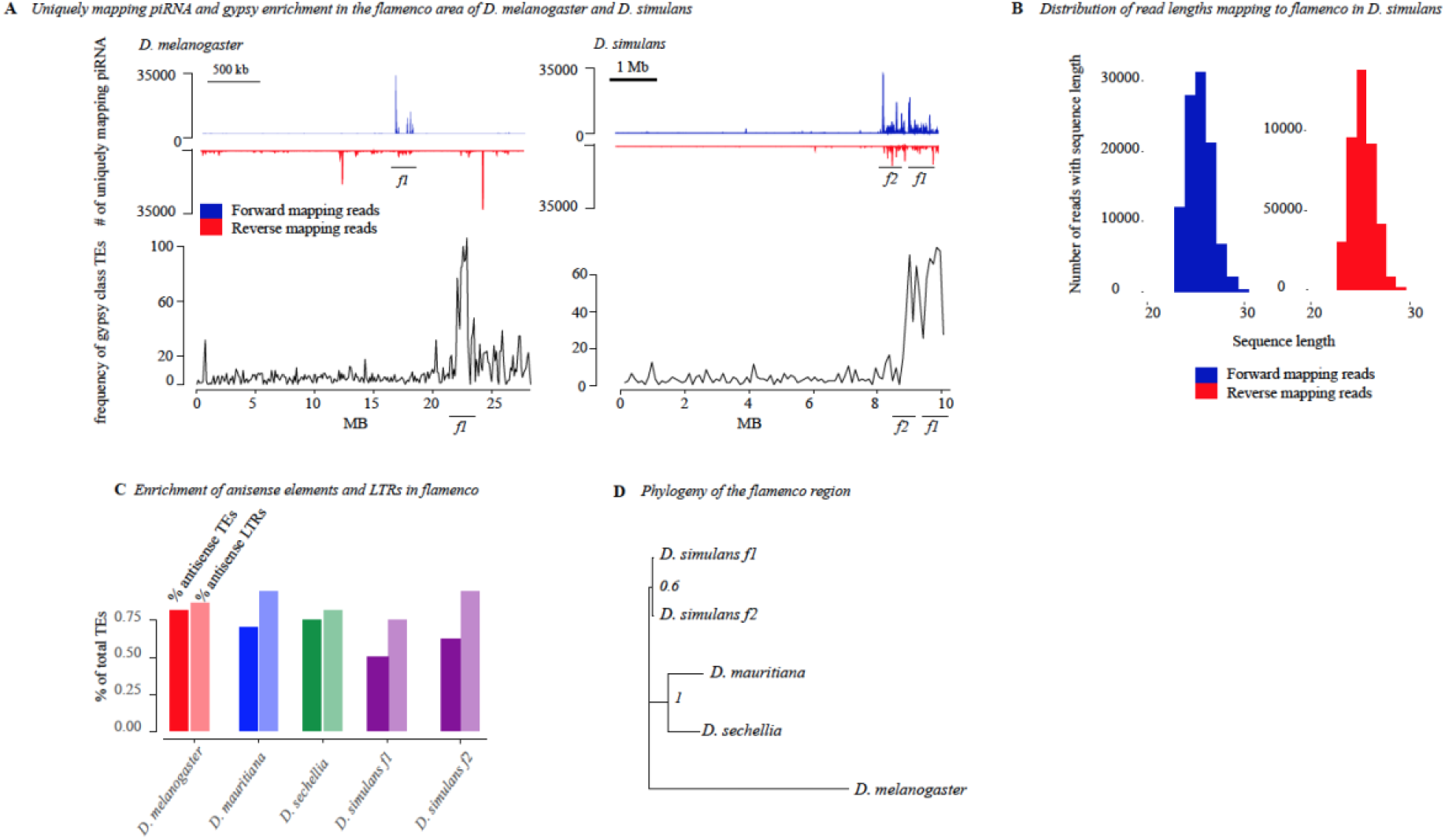
A) Unique piRNA from the ovary and *gypsy* enrichment around *flamenco* and its duplicate in *D. simulans* and *D. melanogaster*. piRNA mapping to the entire contig that contains *flamenco* is shown for both species. The top of the panel shows piRNA mapping to f*lamenco* and is split by antisense (blue) and sense (red) piRNA. The bottom panel shows the frequency of *gypsy*-type transposon annotations across the contig containing *flamenco*, counted in 100 kb windows. There is a clear enrichment of *gypsy* in the area of *flamenco* and, in *D. simulans*, its duplicate compared to the rest of the contig. B) The distribution of read size for small RNA mapping to *flamenco*. The peak is at approximately 26 bp, within the expected range for piRNA. C) The percent of TEs in *flamenco* in each species which are in the antisense orientation (first bar) and the percent of TEs in the antisense orientation that are also LTR class elements (second bar). D) A phylogenetic tree of the *dip1* and *flamenco* enhancer region for *D. melanogaster* and the *simulans* clade. This region is conserved and alignable between all species. The tree was generated with Mr. Bayes (62).

### Structure of the flamenco locus

*D. melanogaster flamenco* bears a characteristic structure, in which the majority of TEs are *gypsy*-class elements in the antisense orientation (79% antisense orientation, 85% of which are *gypsy* elements) (Figure 2C; Supplementary Table 3). In *D. simulans*, *flamenco* has been colonized by large expansions of *R1* transposable element repeats such that on average the percent of antisense TEs is only 50% and the percent of the locus comprised of LTR elements is 55%. However, 76% of antisense insertions are LTR insertions, thus the underlying *flamenco* structure is apparent when the *R1* insertions are disregarded (Figure 2C). In *D. mauritiana flamenco* is 71% antisense, and of those antisense elements it is 85% LTRs. Likewise in *D. sechellia* 78% of elements are antisense, and of those 81% are LTRs. *flamenco* retains the overall structure of a canonical *D. melanogaster*-like *flamenco* locus in all of these species, however in *D. simulans* the nature of the locus is somewhat altered by the abundant *R1* insertions (Figure 2C).

### flamenco is duplicated in D. simulans

In *D. simulans*, we unexpectedly observed that *flamenco* is duplicated on the X chromosome; the duplication was confirmed with PCR and a restriction digest (Figure 1, Supplementary File 2). These duplications are associated with a conserved copy of the putative *flamenco* enhancer as well as copies of the *dip1* gene located proximal to *flamenco* in *D. melanogaster* (Figure 1, 3A). While it is unclear which copy is orthologous to *D. melanogaster flamenco*, all *D. simulans* lines bear one copy that aligns across genotypes. We refer to this copy as *D. simulans flamenco*, and the other copies as duplicates. Otherwise, *flamenco* duplicates do not align with one another and lack synteny amongst their resident TEs. Possible evolutionary scenarios are that the *flamenco* duplication occurred early in the *simulans* lineage, that the clusters evolved very rapidly, or that the duplication encompassed only the promoter region and was subsequently colonized by TEs (Figure 1A, 3A).

**Figure 3.**
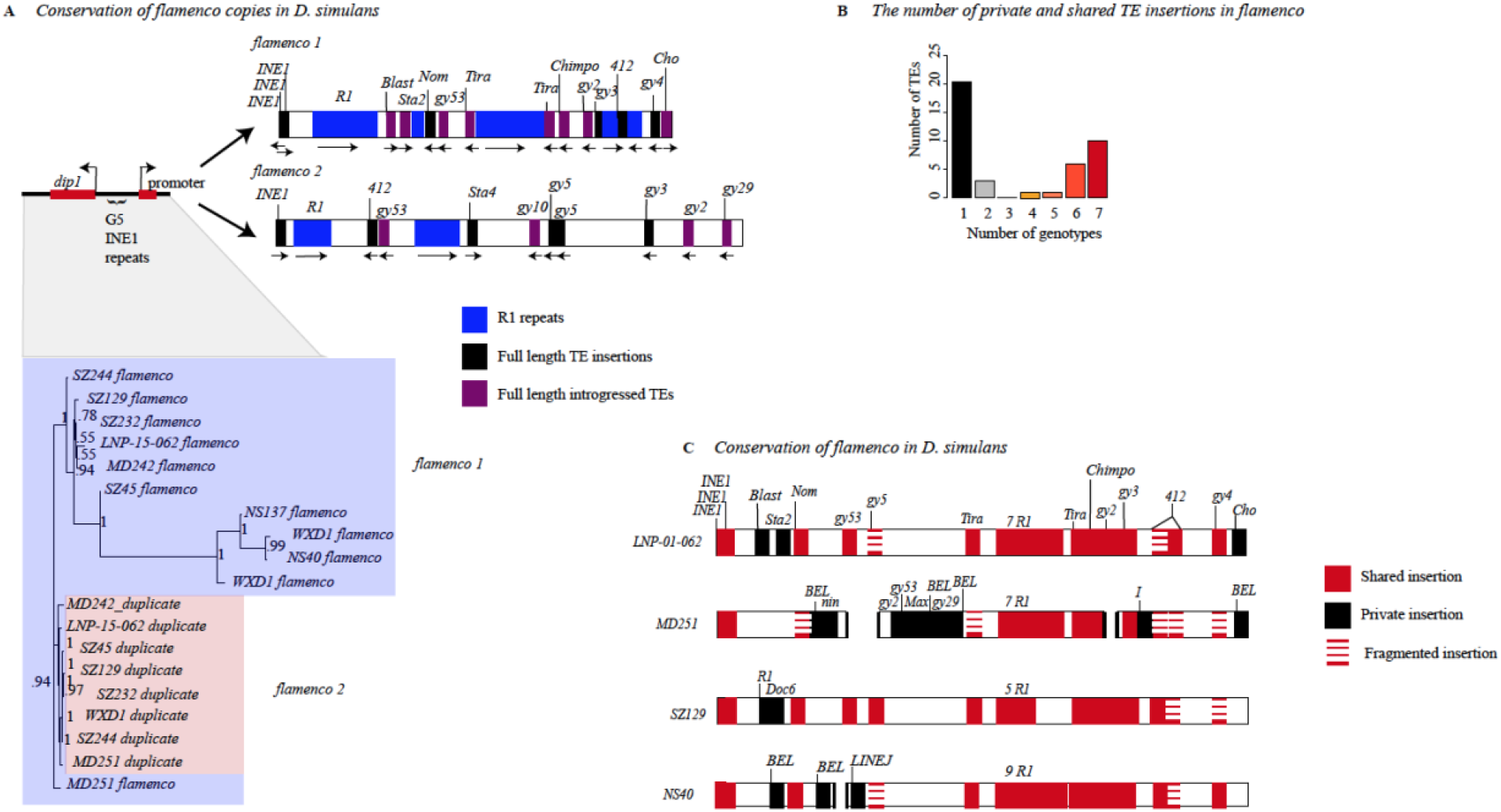
A) Divergence between copies of flamenco. Proximal is a phylogenetic tree of *dip1* and the *flamenco* promoter region from each genome. In between *dip1* and the promoter are a series of *G5/INE1* repeats that are found in every genome. Overall this region is fairly conserved, with the duplicate copies all grouping together with short branch lengths (shown in pink). The original copy of *flamenco* is more diverse with some outliers (shown in light blue) but there is good branch support for all the deep branches of the tree. Distal is a representation of *flamenco* and its duplicate. R1 repeat regions are shown in blue. Full length transposable elements are labeled. There is no synteny conservation between *flamenco* and its duplicate. B) The proportion of insertions that are shared by one through seven genotypes (genotypes with complete *flamenco* assemblies). C) Divergence of flamenco within *D. simulans*. Labeled TEs correspond to elements which are present in a full length copy in at least one genome. If they are shared between genomes they are labeled in red, if they are unique they are black. If they are full length in one genome and degraded in other genomes they are represented by stacked dashes. If they are present in the majority of genomes but missing in one, it is represented as a missing that TE, which is agnostic to whether it is a deletion or the element was never present

The *flamenco* duplicate is absent in the *D. simulans* reference assembly, *w^501^* (GCA_000754195.3), but present in *wxD^1^*, suggesting it was polymorphic or absent between the collection of these strains (or was not assembled). The duplicate retains the structure of *flamenco*, with an average of 67% of TEs in the antisense orientation, and 91% of the TEs in the antisense orientation are LTRs. The duplicate of *flamenco* is less impacted by *R1*, with some genotypes having as few as 8 *R1* insertions (Figure 3C).

### R1 LINE elements at the flamenco locus

*R1* elements are well-known to insert into rDNA genes, are transmitted vertically, and evolve similarly to the genome background rate (28). They have also been found outside of rDNA genes, but only as fragments. *R1* elements are absent from *flamenco* in the *D. simulans* reference assembly, aside from a single fragmented *R1-1* element (*w^501^*). However, as mentioned, *R1* elements are abundant within *flamenco* loci in the *simulans* clade. Outside of *flamenco*, *R1* elements in *D. simulans* are distributed according to expectation, with full length elements occurring only within rDNA (Supplementary File 3). Within *flamenco*, most copies of *R1* occur as tandem duplicates, creating large islands of fragmented *R1* copies (Figure 3A). They are on average 3.7% diverged from the reference R1 from *D. simulans*. Across individual *D. simulans* genomes, ~99 kb of *flamenco* loci consists of *R1* elements, i.e. 26% of their average total length. *SZ45, LNP-15-062, NS40, MD251*, and *MD242* contain 4-7 full length copies of *R1* in the sense orientation, even though all but *SZ45* bear fragmented *R1* copies on the antisense strand. (The *SZ45 flamenco* assembly is incomplete). As the antisense *R1* copies are expected to suppress *R1* transposition, *flamenco* may not suppress these elements effectively. Alternatively, it is possible that *D. simulans flamenco* is still mostly active in the soma, while *R1* is active in the germline, and thus escapes host control by *flamenco*.

In *D. mauritiana*, *flamenco* harbors abundant fragments or copies of *R1* (19 on the reverse strand and 20 on the forward strand), and only one large island of *R1* elements. In total, *D. mauritiana* contains 84 kb of *R1* sequence within *flamenco*. In *D. mauritiana* there are 8 full length copies of *R1* at the *flamenco* locus, 7 in antisense, which are not obviously due to a segmental or local duplication. Finally, we find that *D. sechellia flamenco* lacks full length copies of *R1*, and it contains only 18 KB of *R1* sequence (16 fragments on the reverse strand). Yet, all the copies are on the sense strand, which would not produce fragments that can suppress R1 TEs. Essentially the antisense copies of *R1* in *D. mauritiana* should be suppressing the TE, but we see multiple full length antisense insertions, and *D. sechellia* has no antisense copies, but we see no evidence for recent *R1* insertions. From this it would appear that whatever is controlling the transposition of *R1* lies outside of *flamenco*.

The presence of long sense-strand *R1* elements within *flamenco* is a departure from expectation (18, 28). There is no evidence of an rDNA gene within the *flamenco* locus that would explain the insertion of *R1* elements there, nor is there precedence for the large expansion of *R1* fragments within the locus. Furthermore, the suppression of *R1* transposition does not appear to be controlled by *flamenco*.

### piRNA production from R1

On average *R1* elements within the *flamenco* locus of *D. simulans* produce more piRNA than any other TE within *flamenco* (Supplementary Table 6). *R1* reads mapping to the forward strand constitute an average of 51% of the total piRNAs within the *flamenco* locus from the maternal fraction, ovary, and testis using weighted mapping. The only exception is the ovarian sample from *SZ232* which is a large outlier at only 5%. However reads mapping to the reverse strand account for an average of 84% of the piRNA being produced from the strand in every genotype and tissue – maternal fraction, testis, or ovary. If unique mapping is considered instead of weighted these percentages are reduced by approximately 20%, which is to be expected given that *R1* is present in many repeated copies. Production of piRNA from the reverse strand seems to be correlated with elements inserted in the sense orientation, of which the vast majority are *R1* elements in *D. simulans* (Supplementary Figure 2). The production of large quantities of piRNA cognate to the *R1* element is seemingly pointless – if *R1* only inserts at rDNA genes and are vertically transmitted there is little reason to be producing the majority of piRNA in response to this element.

In *D. sechellia* there are very few piRNA produced from *flamenco* in these tissues, and there are no full length copies of *R1*. Likewise overall weighted piRNA production from *R1* elements on either strand is 2.8-5.9% of the total mapping piRNA. In contrast in *D. mauritiana* there are full length *R1* elements and abundant piRNA production in the maternal fraction and testis. In *D. mauritiana* an average of 28% of piRNAs mapping to the forward strand of *flamenco* are arising from *R1*, and 33% from the reverse strand. In *D. mauritiana R1* elements make up a smaller proportion of the total elements in the sense orientation (24%), versus *D. simulans* (55%).

### Conservation of flamenco

The *dip1* gene and promoter region adjacent to each copy of *flamenco* are very conserved both within and between copies of *flamenco* (Figure 3A). The phylogenetic tree of the area suggests that we are correct in labeling the two copies as the original *flamenco* locus and the duplicate (Figure 3A). The original *flamenco* locus is more diverged amongst copies while the duplicate clusters closely together with short branch lengths (Figure 3A). The promotor region is also conserved and alignable between *D. melanogaster*, *D. sechellia*, *D. mauritiana*, and *D. simulans* (Figure 1D). However, the same is not true of the *flamenco* locus itself. Approximately 3 kb from the promoter *flamenco* diverges amongst genotypes and species and is no longer alignable by traditional sequence-based algorithms, as the TEs are essentially presence/absence polymorphisms that span multiple kb. There is no conservation of *flamenco* between *D. melanogaster*, *D. simulans*, *D. sechellia*, and *D. mauritiana* (Figure 4). However, within the *simulans* clade many of the same TEs occupy the locus, suggesting that they are the current genomic invaders in each of these species (Figure 4).

**Figure 4).**
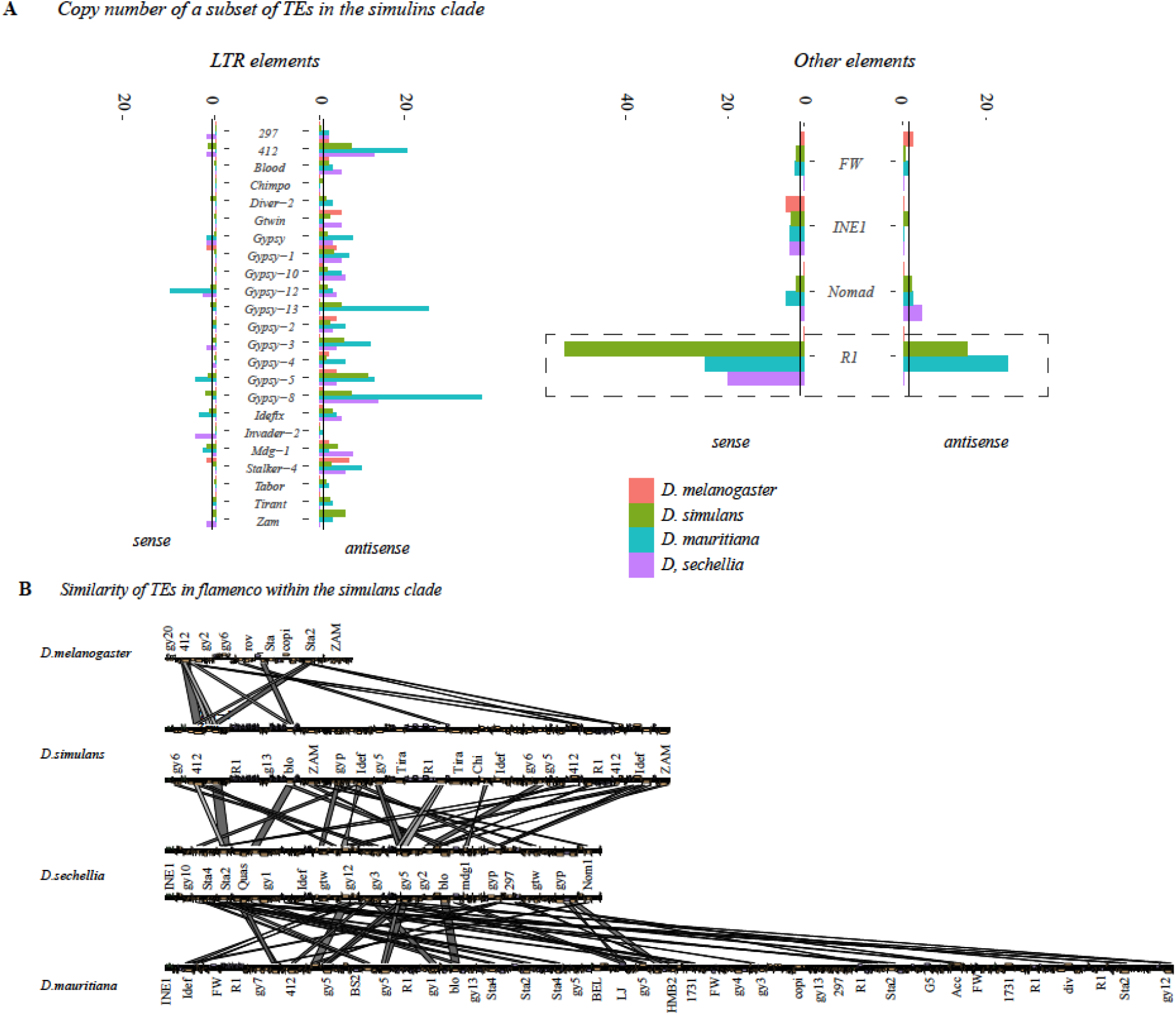
A. Copy number of a subset of transposable elements at *flamenco*. Solo LTRs are indicated by in a lighter shade at the top of the bar. The black line on each bar graph indicates a copy number of one. Values for *D. simulans* are the average for all genotypes with a complete *flamenco* assembly. Note that in *D. melanogaster* (green) most TEs have a low copy number. The expansion of *R1* elements in the *simulans* clade is clearly indicated on the right hand panel with a dotted box. Many elements within *flamenco* are multicopy in the *simulans* clade. While some of this is likely due to local duplications it is clearly a different pattern than *D. melanogaster*. Enrichment of LTR elements on the antisense strand is clear for all species. **B**. Alignment of *flamenco* in *D. melanogaster*, *D. simulans*, *D. sechellia*, and *D. mauritiana*. There is no conserved synteny between species but there are clearly shared TEs, particularly within the *simulans* clade. The expansion of *D. mauritiana* compared to the other species is apparent.

In *D. simulans* the majority of full length TEs are singletons – 54% in *flamenco* and 64% in the duplicate. Copies that are full length in one genotype but fragmented in others are counted as shared, not singletons. Almost half of these singletons in the duplicate are due to a single genotype with a unique section of sequence, in this case *MD251*. Singletons are the single largest category of transposable element insertions, followed by fixed insertions. Thus even within a single population there is considerable diversity at the *flamenco* locus, and subsequently diversity in the ability to suppress transposable elements. For example, *gypsy-29* is present in three genotypes either in *flamenco* or the duplicate, which would suggest that these genotypes are able to suppress this transposable element while the other genotypes are not. In contrast *gypsy-3* is present in more than one full length copy in *flamenco* and its duplicate in every genotype but one where it is present in a single copy. There are a number of these conserved full length TEs that are present in all or nearly all genotypes, including *Chimpo*, *gypsy-2*, *Tirant*, and *gypsy-4*. In addition, the *INE1* elements adjacent to the promoter are conserved.

It is notable that any full length TEs are shared across all genotypes, given that *wxD^1^* was like collected 30-50 years prior to the others, and the collections span continents (Figure 3C). Two facts are relevant to this observation: (1) TEs were shown not to correlate with geography (67) and (2) *D. simulans* is more diverse within populations than between different populations (68–70). Other explanations are also plausible. Selection could be maintaining these full length TEs, *wxD^1^* could have had introgression from other lab strains, or a combination of these explanations.

### Suppression of TEs by the flamenco locus and the trap model of TE control

In *D. melanogaster*, it was proposed that while germline clusters may have many insertions of a single TE, the somatic ‘master regulator’ *flamenco* will have a single insertion of each transposon, after which they are silenced and no longer able to transpose (18).

Here, we evaluate the following lines of evidence to determine if they support the trap model of transposable element suppression. (1) How many TEs have antisense oriented multicopy elements within *flamenco*? (2) How many TEs have full length and fragmented insertions, suggesting the older fragments did not suppress the newer insertion? (3) How many *de novo* insertions of TEs in the *flamenco* duplicate of *D. simulans* are also present in the original *flamenco* copy?

#### How many TEs have antisense oriented multicopy elements within *flamenco*?

Due to the difficulty in classifying degraded elements accurately, for example between multiple classes of *gypsy* element, we will focus here on full length TEs, suggesting recent transposition. In *D. melanogaster* there are 7 full length elements, none of which are present in more than one antisense copy. These elements make up 27% of the *flamenco* locus. Full length copies of five of these elements were also reported previously for other strains of *D. melanogaster* (18)

In *D. sechellia* there are 14 full length TEs within the *flamenco* locus, three of which are present in multiple copies. Two of these, *INE1* and *412*, are likely present due to local duplication. In particular the *INE1* elements flanking the promoter, are in the sense orientation, and are conserved between *D. sechellia, D. mauritiana*, and *D. simulans*. The only element present in multiple antisense copies is *GTWIN*. Similar to *D. melanogaster* these 14 elements make up 27% of the *flamenco* locus.

*D. mauritiana* contains 22 full length TEs within the *flamenco* locus. Four of these are present in multiple antisense full length copies – *INE1*, *R1*, *Stalker-4*, and *Cr1a*. While some of the five antisense copies of *R1* likely originated from local duplications – they are in the same general region and tend to be flanked by *gypsy-8*, not all of them show these patterns. Furthermore, as aforementioned, there also are full length sense copies of *R1* suggesting *R1* is not being suppressed by *flamenco*. *gypsy-12* and *gypsy-3* have a second antisense copy within *flamenco* that is just below the cutoff to be considered full length – in *gypsy-3* the second copy is 10% smaller, for *gypsy-12* it is present but missing an LTR. Full length TEs make up 19% of the *flamenco* locus.

In *D. simulans* there are 24 full length TEs present in any of the seven complete *flamenco* assemblies. Six of these are present in multiple antisense copies within a single genome – *INE1*, *Chimpo*, *gypsy-4*, *412*, *Tirant*, and *BEL-unknown*. The two *Tirant* copies are likely a segmental duplication as they flank an *R1* repeat region. In addition, most *INE1* copies are present proximal to the promoter as aforementioned. *Chimpo* is present in three full length copies within *MD242 flamenco*, with no evidence of local duplication. While there are no full length copies of *R1* inserted in antisense, *R1* is present in full length sense copies despite many genomes containing antisense fragments, suggesting *flamenco* is not suppressing *R1*. On average full length TEs constitute 20% of *flamenco* in *D. simulans*.

In the duplicate of *flamenco* in *D. simulans* there are 30 full length TEs present in any one of the five complete *flamenco* duplicate assemblies. However, none of them are multicopy in antisense. However, they are multicopy relative to the original copy of *flamenco*. *gypsy-3*, *BEL-unknown*, *Nomad-1*, *Chimpo*, *gypsy-53A*, *R1*, and *INE1* are all multicopy with respect to the original *flamenco* within a given genome. Some of these may have been inherited at the time of duplication, however are full length in both copies suggesting recent transposition. In the duplicate of *flamenco* full length TEs occupy an average of 17% of the locus. *MD251* is an exception which weights the average, with 28% of the locus, while between 10 and 15% is found for the remaining copies. Thus *D. simulans* and *D. mauritiana* overall do not meet the expectation that *flamenco* will contain a single insertion of any given TE.

#### How many TEs have full length and fragmented insertions?

Full length elements are younger insertions than fragmented insertions. If a full length element is inserted in *flamenco* and there are fragments in the antisense orientation elsewhere in *flamenco* this indicates that *flamenco* did not successfully suppress the transposition of this element.

In *D. melanogaster* two elements have fragments in antisense and a full length TE – *Doc* and *Stalker-2*. *D. sechellia* has 9 elements that are present as a full length TE and a fragment in antisense (including *412*, *GTWIN*, *mdg-1*, and *nomad*) and 6 that are multicopy that are due to a solo LTR (including *blood*, *297*, and *Stalker-4*). *D. mauritiana* has 21 elements that are present in full length and a fragment in antisense (including *blood*, *412*, *gypsy-10-13*, and *R1*), and four elements that are multicopy due to a solo LTR (*mdg-1*, *Idefix*, and *gypsy-7,10)*.

In *D. simulans*, TEs that fit this criteria in *flamenco* include *gypsy-2*, *gypsy-3*, *gypsy-4*, *gypsy-5*, *Chimpo*, *412*, *INE1*, *R1*, *Tirant*, and *Zam*. *297* and *Nomad-1* are present in full length copies but only multi-copy in the context of solo LTRs. In the duplicate of *flamenco* in *D. simulans* this includes *gypsy-2*, *gypsy-3*, *gypsy-5*, *297*, *Stalker-4*, and *R1*. For example in *NS40* there are 7 full length copies of *R1* in the sense orientation that likely duplicated in place, as well as 12 partial copies in the antisense orientation. In the *simulans* clade either fragments of TEs are not sufficient to suppress transposable elements or some elements are able to transpose despite the hosts efforts to suppress them.

### Is flamenco a trap for TEs entering through horizontal transfer?

High sequence similarity between TEs in different species suggests horizontal transfer (71). However, because sequence similarity can also exist due to vertical transmission we will use sequence similarity between R1 elements (inserted at rDNA genes) as a baseline for differentiating horizontal versus vertical transfer. There has never been any evidence found for horizontal transfer of *R1* and it is thought to evolve at the same rate as nuclear genes in the *melanogaster* subgroup (18, 28). Of the full length elements present in any genome at *flamenco* 62% of them appear to have originated from horizontal transfer. This is similar to previous estimates for *D. melanogaster* in other studies (18). Transfer appears to have occurred primarily between *D. melanogaster*, *D. sechellia*, and *D. willistoni*. This includes some known horizontal transfer events such as *Chimpo* and *Chouto* (72), and others which have not been recorded such as *gypys-29* (*D. willistoni*) and the *Max-element* (*D. sechellia*) (Supplementary File 4). The duplicate of *flamenco* is similar, with 53% of full length TEs originating from horizontal transfer. They are many of the same TEs, with a 46% overlap, thus *flamenco* and its duplicate are trapping many of the same TEs. Both *flamenco* and the duplicate the region appears to serve as a trap for TEs originating from horizontal transfer.

In *D. melanogaster* 85% of full length TEs appear to have arisen through horizontal transfer, which is consistent with previous estimates (18). In *D. sechellia* 53% of full length TEs have arisen from horizontal transfer, including some known to have moved by horizontal transfer such as *GTWIN* (*D. melanogaster*/*D. erecta*) (72). *D. mauritiana* has 68% of its full length TEs showing a closer relationship than expected by vertical descent with TEs from *D. sechellia*, *D. melanogaster*, and *D. simulans*. The hypothesis that *flamenco* serves as a trap for TEs entering the population through horizontal transfer holds throughout the *simulans* clade.

### Flamenco piRNA is expressed in the testis and the maternal fraction

Canonically, *flamenco* piRNA is expressed in the somatic follicular cells of the ovary and not in the germline, and also does not produce a ping pong signal (24). It was not thought to be present in the maternal fraction of piRNAs or other tissues. However, that appears to be variable in different species (Figure 5). We examined single mapping reads in the *flamenco* region from testes and embryos (maternal fraction) in *D. simulans*, *D. mauritiana*, *D. sechellia*, and *D. melanogaster*. As a control we also included *D. melanogaster* ovarian somatic cells, where Aub and Ago3 are not expressed and therefore there should be no ping pong signals. In *D. simulans* and *D. mauritiana flamenco* is expressed bidirectionally in the maternal fraction and the testis, including ping pong signals on both strands (Figure 5A & C; Supplementary Figure 1). In *D. sechellia*, there is no expression of *flamenco* in either of these tissues. Discarding multimappers in the maternal fraction 63% (*D. mauritiana*) – 36% (*D. simulans*) of the ping pong signatures on the X with a z-score of at least 0.9 are located within *flamenco* (Figure 5C). In the testis the picture is more complicated – in *D. mauritiana* 50% of ping pong signals on the X with a z-score of at least 0.9 are located within *flamenco*, which amounts to a substantial ping pong signature (Supplementary Figure 1). While mapping of piRNA to both strands was observed in *D. simulans* testis, there is very little apparent ping pong activity (5 positions in *flamenco* z > 0.9; 15 potential ping pong signals on the X). In *D. melanogaster*, there is uni-strand expression in the maternal fraction, but it is limited to the region close to the promoter. In *D. melanogaster* no ping pong signals have a z-score above 0.8 in the maternal fraction or the ovarian somatic cells. There are ping pong stacks in *flamenco* in the testis of D. melanogaster (2% of the total on the contig), however they are limited to a single region and are not abundant enough to be strong evidence of ping pong activity.

**Figure 5).**
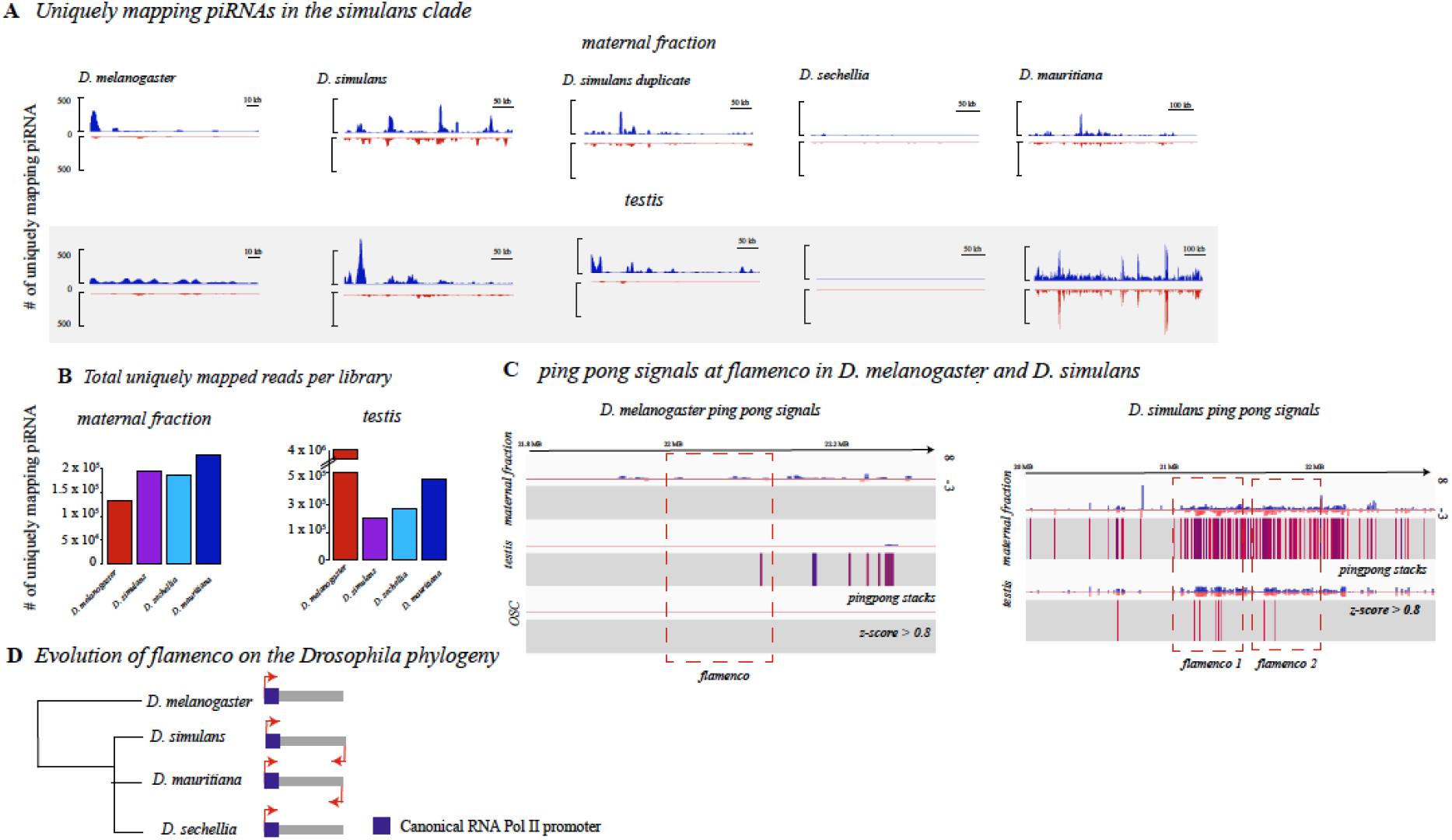
**A**. Expression of single mapping piRNAs in the maternal fraction and testis (gray) of *D. melanogaster* and the *simulans* clade. Antisense mapping reads are shown in blue, sense in red. Libraries are RPM normalized and scaled across library type. *D. sechellia* has no expression of *flamenco* in the maternal fraction or the testis. *D. melanogaster* has low expression in the maternal fraction and very little ping pong activity. *D. simulans* and *D. mauritiana* show dual stranded expression in the testis and maternal fraction. **B**. The total number of uniquely mapping reads for each of the libraries illustrated in A. This is included to demonstrate that a low number of mapping reads does not explain the patterns seen in *D. sechellia* versus *D. mauritiana*. **C**. The height of 10 nt pingpong stacks at *flamenco* in *D. melanogaster* maternal fraction, testis and ovarian somatic cells is shown on the left. Below each schematic of the height of the stacks is the position of z-scores over 0.8, indicating the likelihood that this is a real ping pong signal as opposed to an artifact. Signals move from red to blue as they approach 1. In the testis a few ping pong signals reach this threshold but not enough to indicate convincingly that there is ping pong activity. On the right are the ping pong stacks and z-scores for the maternal fraction and testis in *D. simulans*. Only in the maternal fraction are the density of z-scores over 0.8 convincing enough to indicate an active ping pong cycle in the *flamenco* region. However, the presence of stacks is enriched in testis, thus this may warrant further investigation. *D. mauritiana* also has convincing ping pong signals in this region (Supplementary Figure 1). **D**. A schematic of the evolution of *flamenco* and its mode expression in the *simulans* and *melanogaster* clade.

In the duplicate of *flamenco* in the maternal fraction 15% of the ping pong signals with a z-score above 0.9 on the X are within the *flamenco* duplicate. The *flamenco* duplicate does not have a strong signal of the ping pong pathway in the testis. In addition, *flamenco* in these species has been colonized by full length TEs thought to be active in the germline such as *blood*, *burdock, mdg-3*, *Transpac*, and *Bel* (73, 74). *blood* is also present in *D. melanogaster* in a full length copy while there is no evidence of germline activity for *flamenco* in *D. melanogaster*, though no other putative germline TEs are present. The differences in ping pong signals between species and the presence of germline TEs in *D. simulans* and *D. mauritiana* suggests that the role of *flamenco* in these tissues has evolved between species.

## Discussion

The piRNA pathway is the organisms primary mechanism of transposon suppression. While the piRNA pathway is conserved, the regions of the genome that produce piRNA are labile, particularly in double stranded germline piRNA clusters (9). The necessity of any single cluster for TE suppression in the germline piRNA pathway is unclear, but likely redundant (9). However, *flamenco* is thought to be the master regulator of the somatic support cells of the ovary, preventing *gypsy* elements from hopping into germline cells (18, 21, 23, 24, 75, 76). It is not redundant to other clusters, and insertion of a single element into *flamenco* in *D. melanogaster* is sufficient to initiate silencing. Here we show that the function of *flamenco* appears to have diversified in the *D. simulans* clade, acting in at least some tissues as a germline piRNA cluster.

### Dual stranded expression of flamenco

In this work, we showed that piRNAs of the *flamenco* locus in *D.simulans* and *D. mauritiana* are deposited maternally, align to both strands, and exhibit ping-pong signatures. This is in contrast to *D. melanogaster*, where *flamenco* acts as a uni-strand cluster in the soma (3), our data thus suggest that the *flamenco* locus in *D. simulans* and *D. mauritiana* acts as a dual-strand cluster in the germline. In *D. sechellia* the attributes of *flamenco* uncovered in *D. melanogaster* appear to be conserved – no expression in the maternal fraction and the testis and no ping pong signals. Given that *flamenco* is likely a somatic uni-strand cluster in *D. erecta*, we speculate that the conversion into a germline cluster happened in the *simulans* clade (3). Such a conversion of a cluster between the somatic and the germline piRNA pathway is not unprecedented. For example, a single insertion of a reporter transgene triggered the conversion of the uni-stranded cluster *20A* in *D. melanogaster* into a dual-strand cluster (77).

The role of *flamenco* in *D. simulans* and *D. mauritiana* as the master regulator of piRNA in somatic support cells may still well be true – the promoter region of the *flamenco* cluster is conserved between species and between copies of *flamenco* within species. This suggests that in at least some contexts (or all) the cluster is still serving as a uni-strand cluster transcribed from a traditional RNA Pol II site (15). However it has acquired additional roles, producing dual strand piRNA and ping pong signals, in these two species, in at least the germline. However, in *D. simulans*, the majority of these reverse stranded piRNAs are emerging from the *R1* insertions within *flamenco*. There is no evidence at present that *R1* has undergone an expansion in function in *D. simulans*, thus it is unclear what, if any, functional impact the reverse stranded piRNAs have at the *flamenco* locus.

### Duplication of flamenco in D. simulans

In *D. simulans*, *flamenco* is present in 2 genomic copies, and this duplication is present in all sequenced *D. simulans* lines except the reference strain. The *dip1* gene and putative *flamenco* promoter flanking the duplication also has a high similarity in all sequenced lines (Fig. 2B). This raises the possibility that the duplication of *flamenco* in *D. simulans* was positively selected. Such a duplication may be beneficial as it increases the ability of an organism to rapidly silence TEs. Individuals with large piRNA clusters (or duplicated ones) will accumulate fewer deleterious TE insertions than individuals with small clusters (or non-duplicated ones), and duplicated clusters may therefore confer a selective advantage (78).

### Rapid evolution of piRNA clusters

A previous work showed that dual- and uni-strand clusters evolve rapidly in *Drosophila* (20). In agreement with this work we also found that the *flamenco*-locus is rapidly evolving between and within species (Fig. 1C, 3B). A major open question remains whether this rapid turnover is driven by selection (positive or negative) or an outcome of neutral processes (eg. high TE activity or insertion bias of TEs). These rapid evolutionary changes at the *flamenco* locus, a piRNA master locus, suggest that there is a constant turnover in patterns of piRNA biogenesis that potentially leads to changes in the level of transposition control between individuals in a population.

## Acknowledgements

Thanks to Colin Meiklejohn for providing some of the fly strains used in this manuscript. We would also like to thank Dimitri Petrov and his lab for providing logistical support to BYK. SS would like to thank Jeff Kittilson for assistance in the laboratory. SS would also like to thank C & F & S Emery for insightful commentary on the manuscript.

## Authors’ contributions

SS conceived the study, performed bioinformatics and drafted portions of the manuscript. FW and RK performed bioinformatics and drafted portions of the manuscript. JV contributed data and bioinformatic analysis. ECL drafted portions of the manuscript and provided data. BYK generated and contributed sequence data.

## Availability of data and materials

All data has been made available in the following repositories:

