## Supplementary figures and images for "Rapid evolutionary diversification of the *flamenco* locus across simulans clade *Drosophila* species"

### Supplemental Figure 1

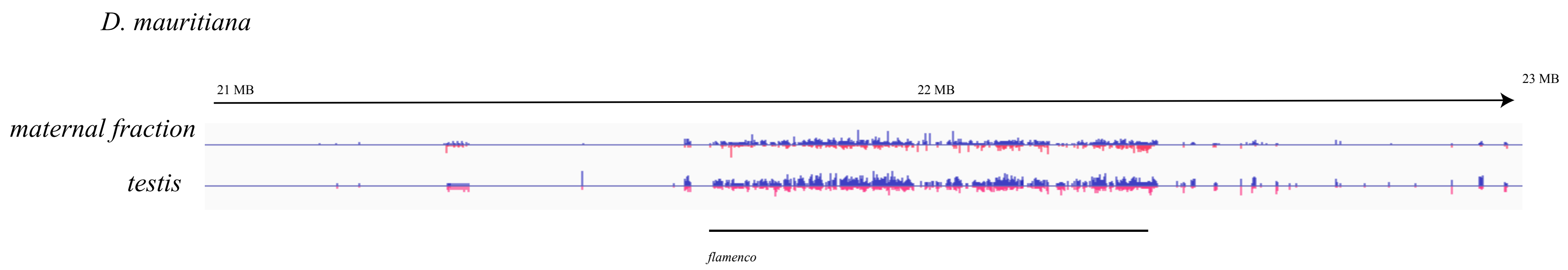

### Supplemental File 1

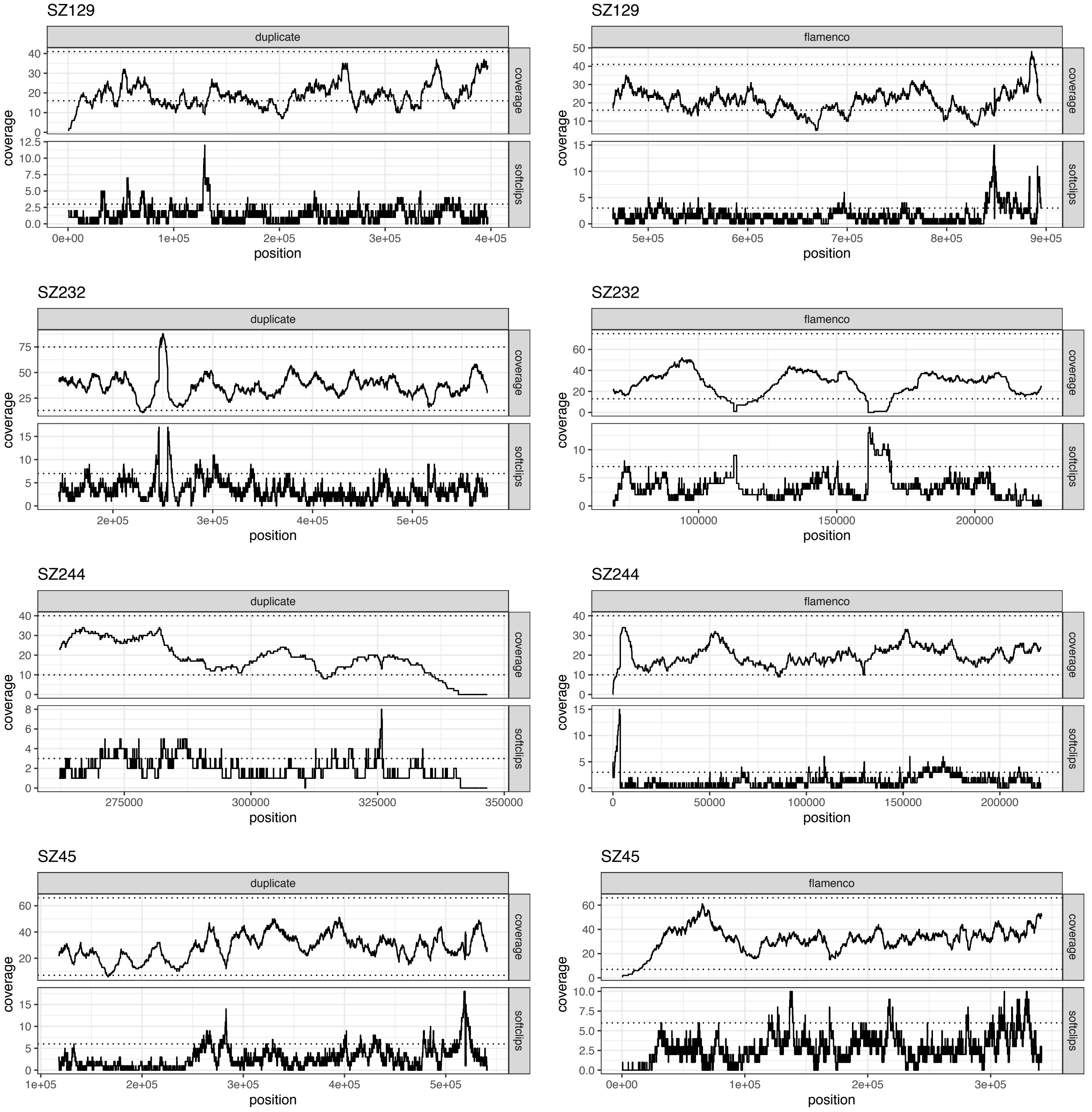

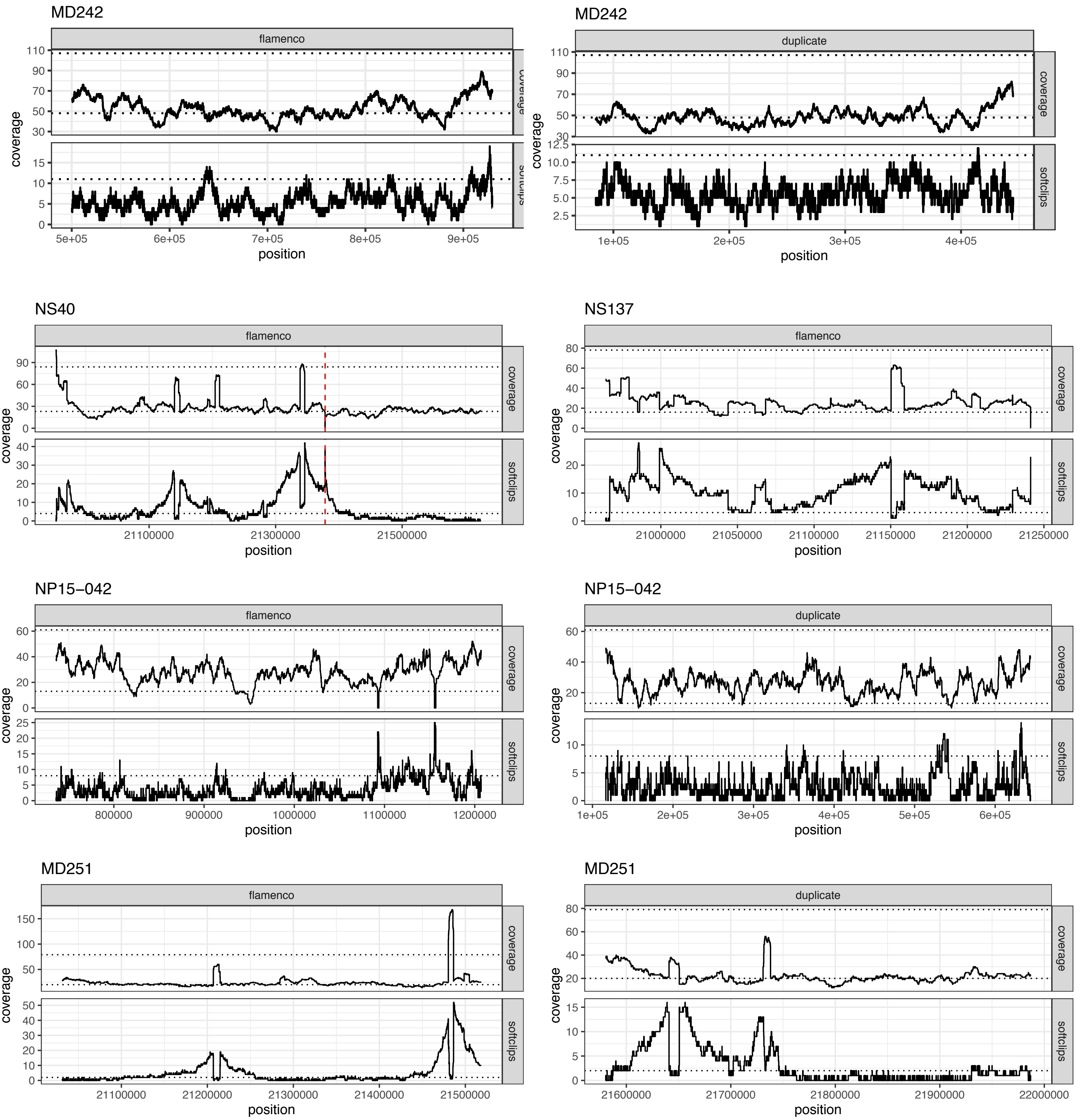

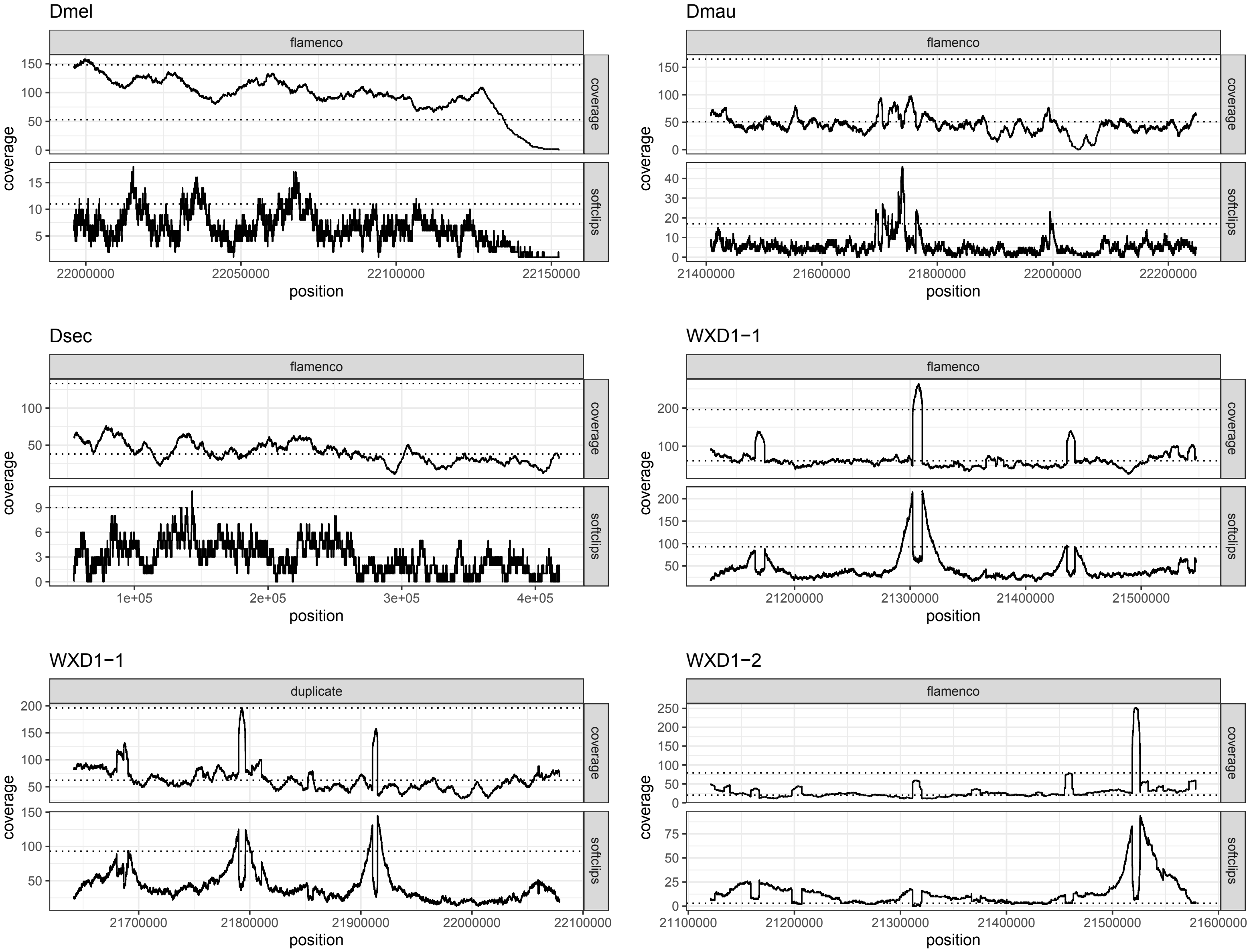
