## Supplemental Figure 2 for "Rapid evolutionary diversification of the *flamenco* locus across simulans clade *Drosophila* species"

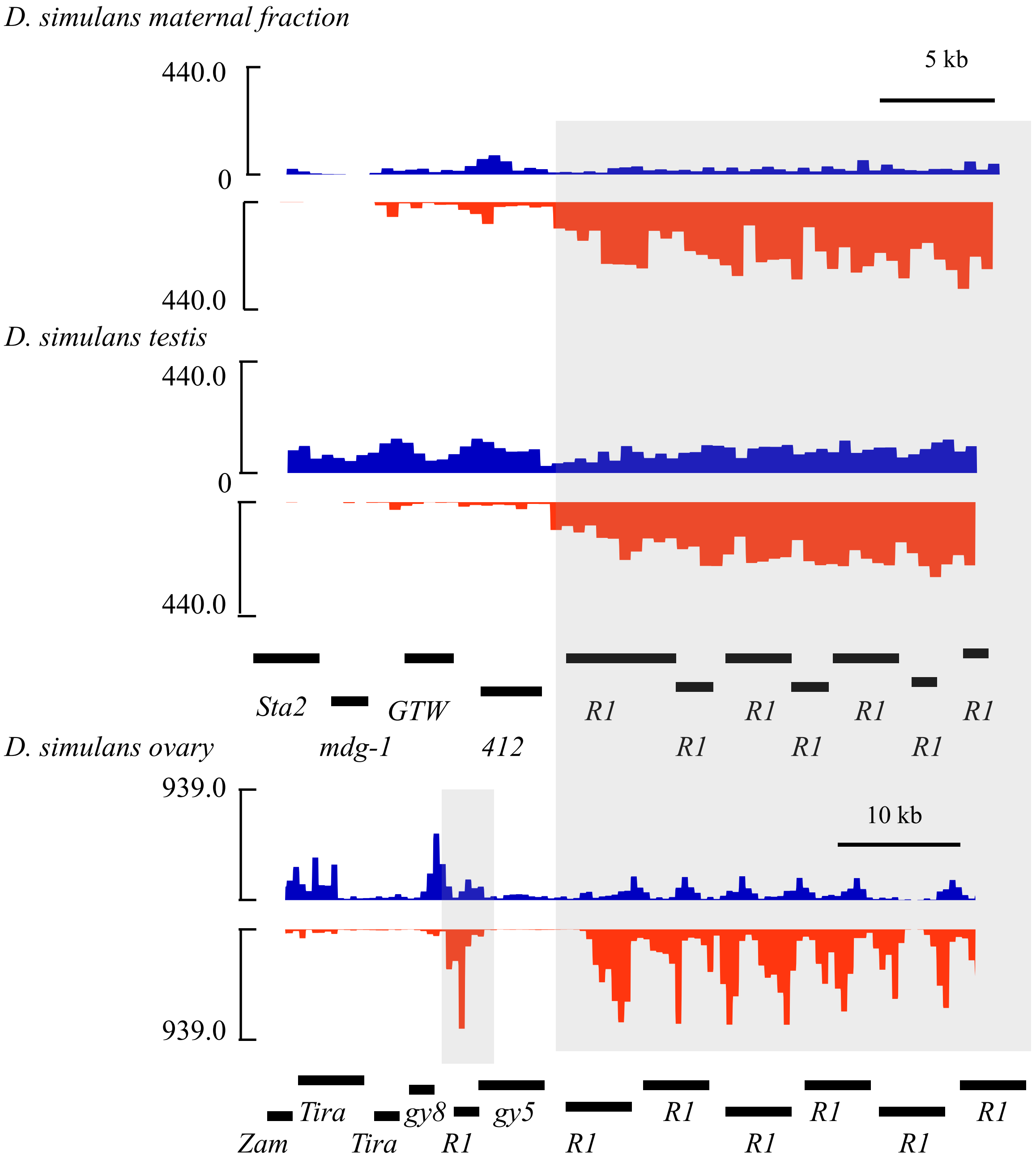


Figure 2: Transcription of piRNA from the reverse strand coordinates with the position of sense orientated R1 elements. Weighted abundance of piRNA mapping is shown for wxD1-1 from the maternal fraction and the testis. piRNA mapping to the forward strand is shown in blue, the reverse strand is shown in red. The bottom panel shows the weighted abundance of piRNA in the ovary for the strain LNP-150-062. Note that this is a different region as the two genotype's flamenco loci do not show large homologous regions.
