## Supplemental Table 1 for "Rapid evolutionary diversification of the *flamenco* locus across simulans clade *Drosophila* species"

| Species/Genotype | # of contigs | N50 | N90 | Assembly Size | Largest Contig |
|---|---|---|---|---|---|
| *SZ 129* | 345 | 21.1 MB | 1.1 MB | 136 MB | 27.6 MB |
| *SZ 232* | 870 | 3.5 MB | 51 KB | 163 MB | 20.3 MB |
| *SZ 244* | 544 | 18.9 MB | 100 KB | 150 MB | 21.2 MB |
| *SZ 45* | 1281 | 2.7 MB | 39 KB | 174 MB | 17.4 MB |
| *LNP-15-062* | 728 | 4.8 MB | 60 KB | 151 MB | 10.5 MB |
| *MD242* | 649 | 8.12 MB | 83 KB | 162 MB | 18.4 MB |
| *MD251* | 25 | 23.5 MB | 430 KB | 136 MB | 28 MB |
| *NS40* | 69 | 21.7 MB | 282 KB | 138 MB | 27 MB |
| *NS137* | 110 | 18.9 MB | 870 KB | 139 MB | 23.3 MB |
| *wxD1-2* | 42 | 13.2 MB | 356 KB | 132 MB | 27 MB |
| *wxD1-1* | 8 | 23.6 MB | 21.7 MB | 129.2 MB | 29 MB |
| *D. melanogaster* | 15 | 28.2 MB | 24.5 MB | 149 MB | 34 MB |
| *D. sechellia* | 378 | 24.8 MB | 131 KB | 153 MB | 28.8 MB |
| *D. mauritianta* | 354 | 24.2 MB | 174 KB | 154 MB | 30.4 MB |
