## Supplemental Table 2 for "Rapid evolutionary diversification of the *flamenco* locus across simulans clade *Drosophila* species"

| Species/Strain | Strain | Chr | Start | End | Identity | Status |
|---|---|---|---|---|---|---|
| *D. mauritiana* | *w12* | X | 21407382 | 22248372 | *flamenco* | complete |
| *D. melanogaster* | *Canton-S* | CM027564.1 | 21996115 | 22152615 | *flamenco* | complete |
| *D. sechellia* | *14021-0248.25* | NW_022611243.1 | 54590 | 418077 | *flamenco* | complete |
| *D. simulans* | *WXD1-1* | X | 21641574 | 22078805 | *duplicate* | complete |
| *D. simulans* | *WXD1-1* | X | 21127235 | 21547722 | *flamenco* | complete |
| *D. simulans* | *WXD1-2* | chr_X | 21120914 | 21579070 | *flamenco* | complete |
| *D. simulans* | *MD251* | chr_X | 21031039 | 21518582 | *flamenco* | complete |
| *D. simulans* | *MD251* | chr_X | 21580275 | 21986909 | *duplicate* | complete |
| *D. simulans* | *SZ45* | tig00001037 | 118032 | 541006 | *duplicate* | complete |
| *D. simulans* | *SZ45* | tig00001107 | 1 | 341500 | *flamenco* | incomplete |
| *D. simulans* | *MD242* | tig00000603 | 500332 | 929454 | *flamenco* | complete |
| *D. simulans* | *MD242* | tig00000603 | 84873 | 444915 | *duplicate* | complete |
| *D. simulans* | *NS137* | chr_X_unlocalized | 20964026 | 21241316 | *flamenco* | complete |
| *D. simulans* | *SZ129* | ig00000300_pilon_pilon | 0 | 397017 | *duplicate* | incomplete |
| *D. simulans* | *SZ129* | tig00000300_pilon_pilon | 464576 | 895002 | *flamenco* | complete |
| *D. simulans* | *SZ232* | tig00000724_pilon_pilon | 69000 | 224020 | *flamenco* | ? |
| *D. simulans* | *SZ232* | tig00000792_pilon_pilon | 146034 | 575012 | *duplicate* | complete |
| *D. simulans* | *SZ232* | tig00000615_pilon_pilon | 6520 | 51023 | *triplicate* | incomplete |
| *D. simulans* | *LNP-15-062* | tig00000577 | 116461 | 644025 | *duplicate* | complete |
| *D. simulans* | *LNP-15-062* | tig00000577 | 736039 | 1206997 | *flamenco* | complete |
| *D. simulans* | *NS40* | chr_X | 20953052 | 21624701 | *flamenco* | complete |
| *D. simulans* | *SZ244* | tig00000001_pilon_pilon | 0 | 221382 | *flamenco* | incomplete |
| *D. simulans* | *SZ244* | tig00000096_pilon_pilon | 262120 | 346650 | *duplicate* | incomplete |
