## Supplemental Table 3 for "Rapid evolutionary diversification of the *flamenco* locus across simulans clade *Drosophila* species"

| Genotype | Identity | % antisense | % of LTR on antisense |
|---|---|---|---|
| *D. mauritiana* | *flamenco* | 0.71 | 0.85 |
| *D. melanogaster* | *flamenco* | 0.78 | 0.82 |
| *D. sechellia* | *flamenco* | 0.78 | 0.81 |
| *wxD1-1* | *flamenco* | 0.55 | 0.82 |
| *wxD1-2* | *flamenco* | 0.56 | 0.77 |
| *wxD1-2* | *duplicate* | 0.57 | 0.85 |
| *LNP-15-062* | *flamenco* | 0.55 | 0.86 |
| *LNP-15-062* | *duplicate* | 0.62 | 0.94 |
| *MD242* | *flamenco* | 0.59 | 0.85 |
| *MD242* | *duplicate* | 0.72 | 0.98 |
| *MD251* | *flamenco* | 0.54 | 0.79 |
| *MD251* | *duplicate* | 0.63 | 0.78 |
| *NS137* | *flamenco* | 0.51 | 0.88 |
| *NS40* | *flamenco* | 0.63 | 0.77 |
| *SZ232* | *flamenco* | 0.51 | 1.00 |
| *SZ232* | *duplicate* | 0.74 | 0.96 |
| *SZ45* | *duplicate* | 0.74 | 0.95 |
| *SZ45* | *flamenco* | 0.60 | 0.88 |
| *SZ129* | *flamenco* | 0.54 | 0.84 |
| *SZ129* | *duplicate* | 0.73 | 1.00 |
